# Gut Competition Dynamics of Live Bacterial Therapeutics Are Shaped by Microbiome Complexity, Diet, and Therapeutic Transgenes

**DOI:** 10.1101/2025.01.21.634159

**Authors:** Nicole Siguenza, Sharyl Bailey, Mohammad Sadegi, Hanna Gootin, Maria Tiu, Jeffrey D. Price, Amanda Ramer-Tait, Amir Zarrinpar

## Abstract

Competitive exclusion is conventionally believed to prevent the establishment of a secondary strain of the same bacterial species in the gut microbiome, raising concerns for the deployment of live bacterial therapeutics (LBTs), especially if the bacterial chassis is a strain native to the gut. In this study, we investigated factors influencing competition dynamics in the murine gut using isogenic native *Escherichia coli* strains. We found that competition outcomes are context-dependent, modulated by microbiome complexity, LBT transgene expression, intestinal inflammation, and host diet. Furthermore, we demonstrated that native LBTs can establish long-term engraftment in the gut alongside a parental strain, with transgene-associated fitness effects influencing competition. We identified various interventions, including strategic dosing and dietary modulation, that significantly enhanced LBT colonization levels by 2 to 3 orders of magnitude. These insights provide a framework for optimizing LBT engraftment and efficacy, supporting their potential translation for human therapeutic applications.

## INTRODUCTION

Live bacterial therapeutics (LBTs) offer unique advantages over traditional pharmaceuticals^1–4^, including the ability to sense and respond to the local environment^5,6^, protect and deliver labile biologics *in situ*^7^, and provide long-term, functionally-curative treatment options^8^ for chronically ill patients. While microbiome-based therapies derived from donor communities^9^ or anaerobic spores^10^ have recently entered the market, engineered LBTs have yet to achieve clinical success. Most LBTs are developed in probiotic bacterial chassis classified as “generally recognized as safe” by the FDA^11^, but these chassis have largely failed to achieve clinical efficacy^12,13^. One prevailing explanation is poor bacterial viability of probiotics as they travel to the intestines. However, researchers have hesitated to adopt more robust native bacteria as chassis, worrying that competitive exclusion will prevent engraftment and long-term therapeutic effects.

The principle of competitive exclusion postulates that two species occupying overlapping ecological niches cannot coexist^14–16^. In microbial systems, this concept has been extended to suggest that closely related bacteria, such as two strains of the same species, are unlikely to co-colonize in the same intestinal environment^17–19^. Although studies have both supported and challenged this generalization^20,21^, concerns over competitive exclusions have deterred the use of native isolates for LBT development. Importantly, existing studies on microbial competitive exclusion have primarily been tested in gnotobiotic models or with a simple synthetic consortia^17,22–24^, leaving the dynamics of isogenic bacterial competition in a complex, intact microbiome largely unexplored.

Beyond competitive exclusion, other factors such as the metabolic burden of engineered functions may reduce LBT fitness relative to parental strains, further complicating engraftment. However, this oversimplified understanding ignores a myriad of factors potentially influencing LBT colonization: surrounding microbiome community^25,26^, transgene function, luminal substrates, and disease pathophysiology. Addressing these variables is critical to advancing LBT development and will provide a better understanding of the role of the microbiome in health and disease.

Here we present an extensive investigation into the dynamics of bacterial competition and LBT engraftment within a conventional murine gut microbiome. Using a native *Escherica coli* isolate from a conventionally-raised C57Bl/6 mouse, we engineered platform strain EcAZ-2^27^ expressing kanamycin resistance and green fluorescent protein^8^. Two LBT strains were developed: one expressing prokaryotic bile salt hydrolase (BSH) that affects host metabolic homeostasis, and another expressing mammalian anti-inflammatory cytokine interleukin-10 (IL10)^8^. For competition studies, a spectinomycin resistant marker was also integrated into the parental strain’s genome. Collectively, our findings identify key variables and interventions that can modulate LBT engraftment and highlight critical considerations for the development and testing of microbiome-based therapies.

## RESULTS

### Arrival order and gut microbiome complexity shape competitive colonization dynamics

The application of native isolates as LBT chassis is limited by concerns over competitive exclusion. Contrary to these concerns, we previously developed a novel native *Escherichia coli* chassis capable of long-term engraftment in conventionally-raised, wildtype hosts^8,27^. This chassis was derived from a specific-pathogen free (SPF) C57BL/6 mouse and engineered to express green fluorescent protein and kanamycin resistance, creating platform strain EcAZ-2. To produce the therapeutic strain EcAZ-2^BSH, CmR^ (formerly referred to as EcAZ-2^BSH+^)^8^, we integrated the bile salt hydrolase (BSH) gene from *Ligilactobacillus salivarius*^28^ and a chloramphenicol resistance gene *cat* into the genome. EcAZ-2^BSH, CmR^ was chosen as our representative LBT in this study due to its glucose-regulating effects^8^. Although we previously demonstrated the robust engraftment capability of EcAZ-2 without pre-screening for competing native *E. coli*, it remained unclear whether this success was due to an absence of competition or inherent competitive fitness. To address this, we directly tested competition between EcAZ-2-based LBTs and a modified parental strain, EcAZ-2^SpecR^, with a spectinomycin resistance gene *aadA1* incorporated into its genome (**Fig. S1A**).

*In vitro* growth assays in rich media (Super Optimal Broth; SOB) revealed no decrease in fitness of EcAZ-2^BSH, CmR^ compared to EcAZ-2 or EcAZ-2^SpecR^, indicating minimal metabolic burden from BSH expression (**Fig. S1B-D**). *In vitro* indirect competition assays using the Cerillo Duet system determined the EcAZ-2^BSH, CmR^ maintained fitness levels comparable to EcAZ-2^SpecR^ (**Fig. 1A-1B**; **Fig. S1E, F**; one-way ANOVA n.s.). This suggests that the engineered modifications did not affect resource acquisition. With fitness preserved *in vitro*, we next explored how these findings translated to competition against a parental strain in more complex conditions *in vivo*.

**Figure 1.**
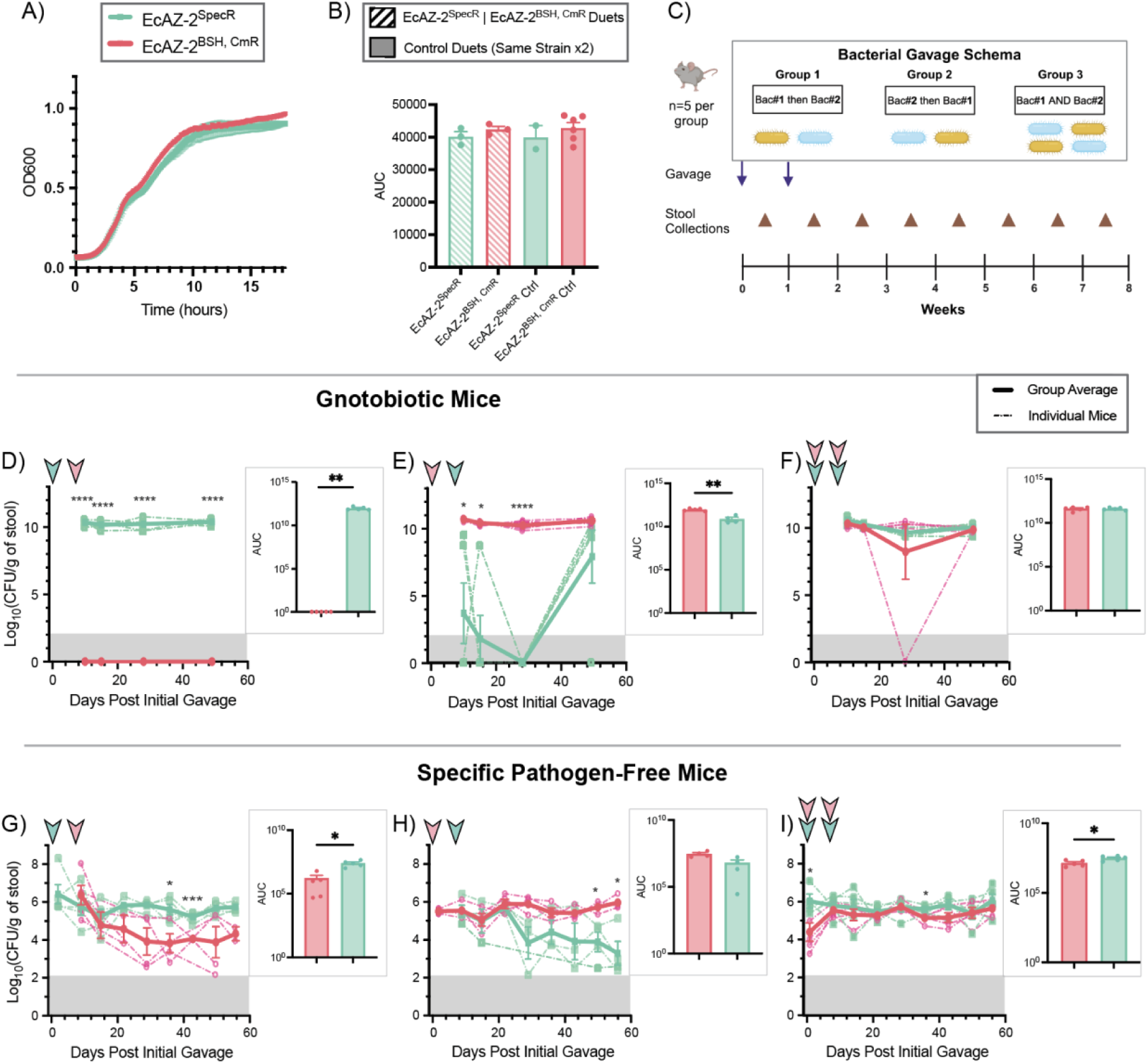
Arrival order and gut microbiome complexity shape competitive colonization dynamics. **(a)** Optical density of bacterial growth in Cerillo Duets. **(b)** AUC of Duet optical density. **(c)** Experimental design to test competition between two bacterial strains in the murine gut. Arrows indicate when the two oral gavages occur. Brown triangles represent time points for stool collections. **(d-f)** Colonization curves for EcAZ-2^BSH, CmR^ (pink) and EcAZ-2^SpecR^ (green) competition in gnotobiotic mice. Arrowheads indicate when each strain was gavaged. Gray area indicates limit of detection. **(g-i)** Colonization curves for EcAZ-2^BSH, CmR^ (pink) and EcAZ-2^SpecR^ (green) competition in specific-pathogen-free mice with intact gut microbiomes. AUC was calculated and compared across strains. Values are represented as mean ± the standard error of mean. Time points with asterisks along the colonization curves indicate significantly different levels in colonization between the two strains as determined by **(d-f)** a Mixed Effects Model or **(g-i)** a two-way ANOVA with a post-hoc Fisher’s LSD test. AUC comparisons were done via two-tailed paired t-test (* p < 0.05; ** p < 0.01; *** p<0.001; no asterisks = not significant).

To evaluate EcAZ-2^BSH, CmR^ competition with EcAZ-2^SpecR^ *in vivo*, a common experimental design was followed: three groups of mice received two oral gavages of ∼10^10^ colony forming units (CFU) of bacteria, spaced one week apart. We administered the bacterial strains in one of three orders: the LBT at t = 0 days and parental strain at t = 7 days; parental strain at t = 0 then LBT at t = 7; or a 50/50 mixture of the LBT and parental strains at t = 0 and t = 7 (**Fig. 1C**). We first tested the co-colonization of EcAZ-2-based strains in germ-free mice. Contrary to the equivalent co-colonization seen for laboratory *E. coli* strain JM109^17^, we found that introducing EcAZ-2^SpecR^ first allowed it to competitively exclude EcAZ-2^BSH, CmR^, leading to the clearance of the LBT from the gut (**Fig. 1D**; two-way ANOVA p < 0.0001 for strains; paired t test of AUC p = 0.0075). Similarly, introduction of EcAZ-2^BSH, CmR^ first results in competition with EcAZ-2^SpecR^ (**Fig. 1E**; two-way ANOVA p < 0.0001 for strains; paired t test of AUC p = 0.0033). However, EcAZ-2^SpecR^ is not cleared from the gut and instead rebounds in colonization levels around 6 weeks post-introduction. Nonetheless, if both bacteria are introduced together, they successfully co-colonize at equivalent levels (**Fig. 1F**; two-way ANOVA n.s.; paired t test of AUC n.s.). These results illustrate the slight fitness advantage EcAZ-2^SpecR^ has over EcAZ-2^BSH, CmR^, and the slight burden of an added transgene, particularly when the parental strain is already established, in gnotobiotic mice.

After determining that EcAZ-2 competes with isogenic strains in gnotobiotic mice, we tested whether these dynamics are recapitulated in an intact murine gut microbiome. Experiments in SPF C57BL/6 mice, without depleted microbiomes, revealed that pre-engraftment of EcAZ-2^SpecR^ does result in competition with EcAZ-2^BSH, CmR^ (**Fig. 1G**; Mixed Effects Model p = 0.0071 for strains; paired t test of AUC p = 0.0244) but the LBT is not excluded from the gut as it had been in gnotobiotic mice (**Fig. 1D**). Instead, EcAZ-2^BSH, CmR^ established colonization at a level that is about 1.5 orders of magnitude lower than that of EcAZ-2^SpecR^ (**Fig. 1G**). Likewise, engrafting EcAZ-2^BSH, CmR^ first results in EcAZ-2^SpecR^ co-colonization at about 2 orders of magnitude lower levels (**Fig. 1H**; Mixed Effects Model p = 0.0351 for strains; paired t test of AUC n.s.). These findings indicate that EcAZ-2^SpecR^ exhibits slightly higher fitness than EcAZ-2^BSH, CmR^, as illustrated by differences in colonization when the bacteria were co-administered (**Fig. 1I**; Mixed Effects Model p = 0.017 for strains; paired t test of AUC p = 0.0479). Nonetheless, EcAZ-2^BSH, CmR^ is able to maintain its transgenic function for the duration of the study (**Fig. S1G**; **Supplemental Table 1**). Overall, arrival order plays a notable role in determining engraftment patterns, while bacterial fitness influences final colonization levels of the individual strains. Furthermore, these results suggest that competition outcomes observed in gnotobiotic mice may only approximate those in a complex microbiome and thus may not be the best indicator of therapeutic potential of an LBT in fully conventional hosts^29–32^.

### Transgene-specific effects modulate competitive fitness

To explore how different therapeutic functions may affect gut competition dynamics, we used a second LBT strain, EcAZ-2^IL10, CmR^ (formerly known as EcAZ-2^IL10+^), that expresses mammalian interleukin-10 (IL-10) and was described in our previous work^8^. Initial *in vitro* growth assays showed no significant fitness deficits in EcAZ-2^IL10, CmR^, suggesting minimal metabolic burden from IL-10 expression (**Fig. S2A**). However, in *in vitro* competition assays using the Duet system, EcAZ-2^IL10, CmR^ exhibited reduced growth when competing against EcAZ-2^SpecR^ (**Fig. 2A**). Specifically, EcAZ-2^IL10, CmR^ displays a prolonged lag phase and slower exponential growth, failing to reach a clear stationary phase (**Fig. 2A,B**; one-way ANOVA p < 0.0001; paired t test of AUC for EcAZ-2^IL10, CmR^ p = 0.0041). Control assays with the same strain inoculated on both sides of the permeable membrane confirmed that decreased fitness of EcAZ-2^IL10, CmR^ was specific to competition with EcAZ-2^SpecR^ (**Fig. S1E; Fig. S2B**). In summation, IL-10 expression may impose a metabolic burden, reducing the LBT’s competitiveness under resource-limited conditions and emphasizing the need to consider transgene-specific effects when designing LBTs.

**Figure 2.**
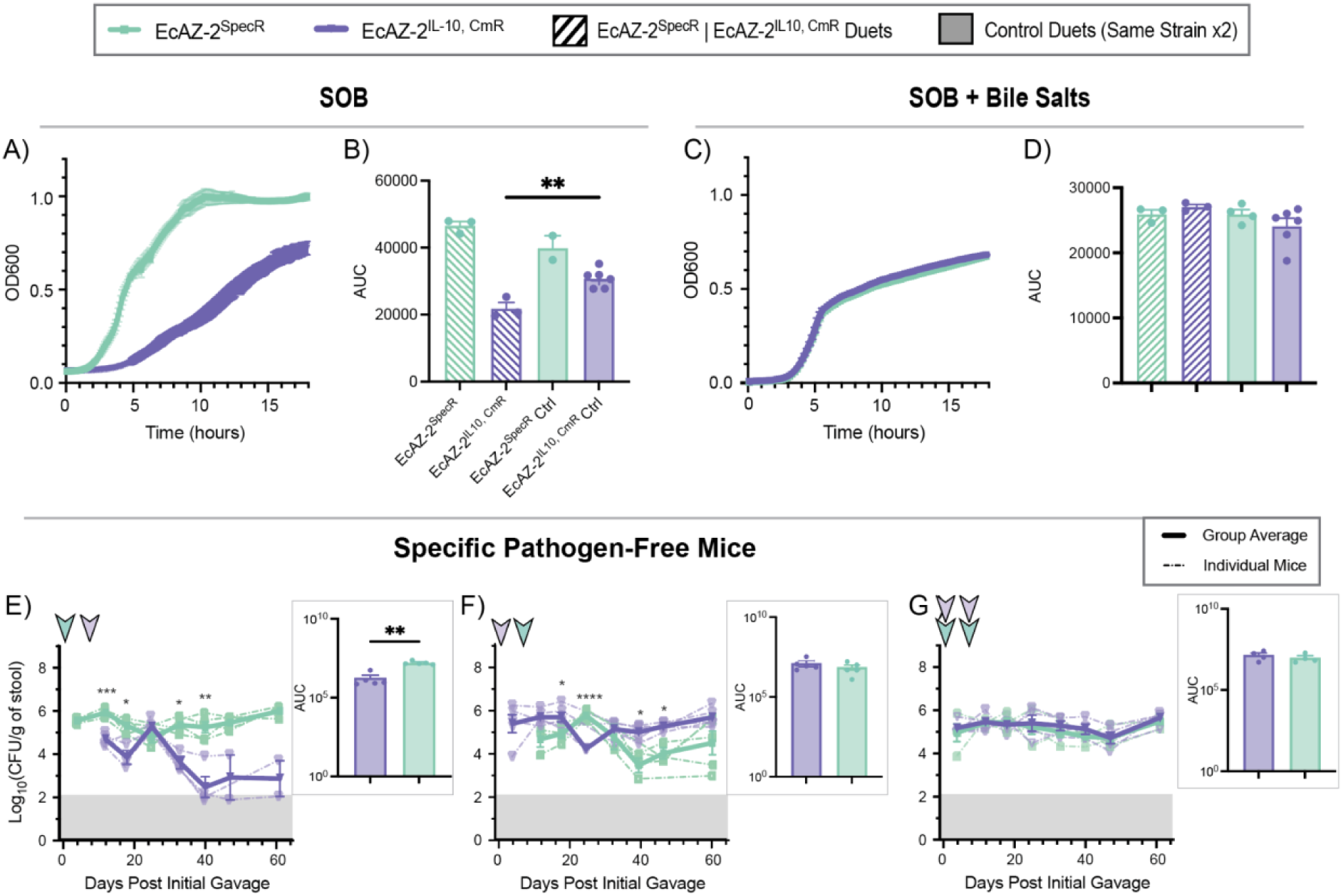
Transgene-specific effects modulate competitive fitness. **(a)** Optical density of bacterial growth in Cerillo Duets using SOB and **(b)** AUC of Duet optical density. **(c)** Optical density of bacterial growth in Cerillo Duets using SOB with bile salts and **(d)** AUC of Duet optical density. **(e-g)** Colonization curves for EcAZ-2^IL10, CmR^ (purple) and EcAZ-2^SpecR^ (green) competition in SPF mice with intact gut microbiome. Arrowheads indicate when each strain was gavaged. Gray area indicates limit of detection. AUC was calculated and compared across strains. Values are represented as mean ± the standard error of mean. Time points with asterisks along the colonization curves indicate significantly different levels in colonization between the two strains as determined by a Mixed Effects Model with a post-hoc Fisher’s LSD test. AUC comparisons were done via two-tailed paired t-test (* p < 0.05; ** p < 0.01; *** p<0.001; no asterisks = not significant).

To determine if luminal substrates such as bile salts could impact the *in vitro* fitness of EcAZ-2^IL10, CmR^, we repeated the competition assays with bile salts added to the medium. This resulted in near identical growth of EcAZ-2^SpecR^ and EcAZ-2^IL10, CmR^. Bile salts depressed the maximum density of stationary phase for both strains but allowed for EcAZ-2^IL10, CmR^ to achieve equivalent growth dynamics with EcAZ-2^SpecR^ (**Fig. 2C, D**; one-way ANOVA n.s.; **Fig. S2C, D**). Likewise, EcAZ-2^BSH, CmR^ and EcAZ-2^SpecR^ exhibited identical growth dynamics with decreased maximum growth when bile salts were added to the Duet *in vitro* competition experiment (**Fig. S2E-G**). This suggests that the suppressive effects of host factors such as bile salts can overshadow the effects of therapeutic burden, highlighting the importance of testing LBT performance under different physiological conditions.

Next, we conducted *in vivo* competition experiments in SPF mice, following the administration schema previously described (**Fig. 1C**). As observed for EcAZ-2^BSH, CmR^, arrival order influenced engraftment patterns but the introduction of a different transgene altered final colonization levels. Pre-engraftment with EcAZ-2^SpecR^ caused EcAZ-2^IL10, CmR^ to engraft at levels 3 orders of magnitude lower (**Fig. 2E**; Mixed Effects Model p < 0.0001 for strains; paired t test of AUC p = 0.0029), as compared to only a 1.5 order of magnitude disparity for EcAZ-2^BSH, CmR^ (**Fig. 1G**). When EcAZ-2^IL10, CmR^ was administered first, the parental strain engrafted at about 1 log_10_-fold lower levels (**Fig. 2F**; Mixed Effects Model p = 0.0099 for strains; paired t test of AUC n.s.), rather than the roughly 2 orders of magnitude by the BSH-expressing LBT (**Fig. 1H**). This difference between LBTs aligns with the metabolic burden detected *in vitro* for EcAZ-2^IL10, CmR^ (**Fig. 2A,B**), potentially reducing the IL10-expressing LBT’s ability to efficiently outcompete EcAZ-2^SpecR^. Importantly, the metabolic burden did not induce function loss over the duration of the study (**Fig. S2H,I**). Finally, co-administration of both strains resulted in equivalent co-colonization (**Fig. 2G**; two-way ANOVA n.s.; paired t test of AUC n.s.), consistent with the *in vitro* competition outcomes observed in the presence of bile salts (**Fig. 2C,D**). Overall, our findings show that introduction of a different transgene with an increased metabolic burden alters competition outcomes both *in vitro* and *in vivo*, particularly in a setting where the bacteria has to compete with an engrafted parental strain.

### Diet-driven effects on bacterial competition outcomes

To explore if *in vivo* alteration of the luminal nutritional profile could alter competition outcomes, we placed mice on high fat diet (HFD) or atherogenic diet (AGD), which is particularly high in cholesterol content, both of which are known to increase fecal bile acid levels^33–36^. Mice were acclimated to their respective diet for 2 weeks prior to gavage (**Fig. 3A**). We chose EcAZ-2^BSH, CmR^ as our representative LBT for these dietary challenges due to its potential therapeutic function of affecting glucose homeostasis by deconjugating bile salts^8^. Moreover, dietary alterations allow us to explore how variations in human diet could affect LBT colonization under diverse host conditions.

**Figure 3.**
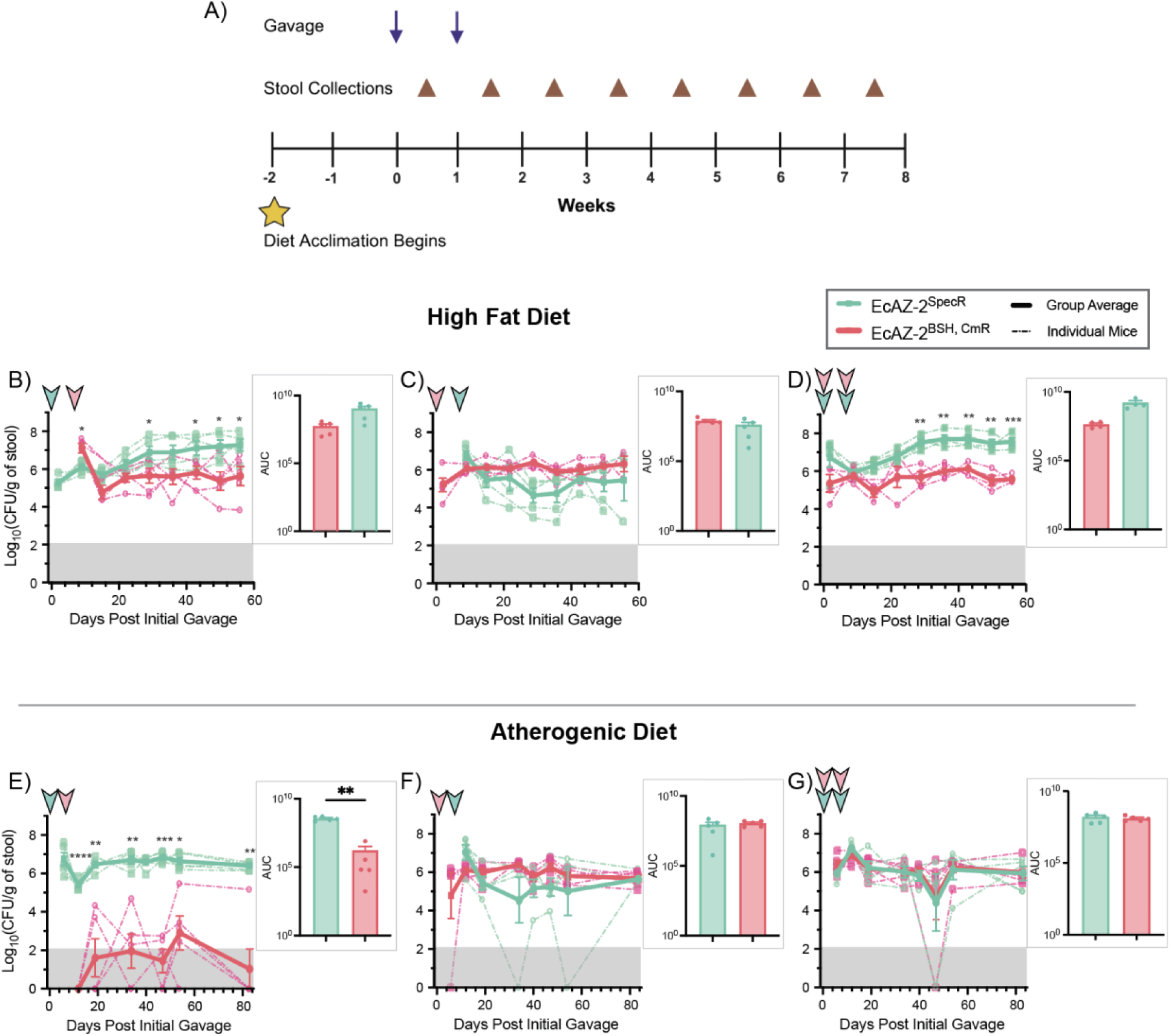
Diet-driven effects on bacterial competition outcomes. **(a)** Experimental design to test competition between two bacterial strains under different dietary conditions. Arrows indicate when the two oral gavages occur. Brown triangles represent time points for stool collections. Yellow star denotes when the mice were started on their respective diets. **(b-d)** Colonization curves for EcAZ-2^BSH, CmR^ (pink) and EcAZ-2^SpecR^ (green) competition in SPF mice consuming HFD. **(e-g)** Colonization curves for EcAZ-2^BSH, CmR^ (pink) and EcAZ-2^SpecR^ (green) competition in SPF mice consuming AGD. Arrowheads indicate when each strain was gavaged. Gray area indicates limit of detection. AUC was calculated and compared across strains. Values are represented as mean ± the standard error of mean. Time points with asterisks along the colonization curves indicate significantly different levels in colonization between the two strains as determined by a Mixed Effects Model with a post-hoc Fisher’s LSD test. AUC comparisons were done via two-tailed paired t-test (* p < 0.05; ** p < 0.01; *** p<0.001; no asterisks = not significant).

Mice on HFD receiving EcAZ-2^SpecR^ first exhibited an engraftment pattern similar to those on normal chow (**Fig. 1G**), where EcAZ-2^BSH, CmR^ engrafts at about 1.5 orders of magnitude lower than the parental strain (**Fig. 3B**; Mixed Effects Model p = 0.011 for strains; paired t test of AUC n.s.). However, HFD allowed for equivalent co-colonization of the two strains when EcAZ-2^BSH, CmR^ engrafted first (**Fig. 3C**; Mixed Effects Model n.s.; paired t test of AUC n.s.). Strikingly, co-administration led to EcAZ-2^SpecR^ engrafting at 2 log_10_-fold higher levels than EcAZ-2^BSH, CmR^ (**Fig. 3D**; Mixed Effects Model p < 0.0001 for strains; paired t test of AUC n.s.), contrasting the equivalent co-colonization previously seen on normal chow (**Fig. 1I**). Overall, HFD diminished the impact of arrival order and bolstered the competitive advantage of the parental strain.

Under AGD, secondary introduction of EcAZ-2^BSH, CmR^ led to diminished colonization and subsequent clearance of the LBT from 4 out of 5 mice (**Fig. 3E**; two-way ANOVA p < 0.0001 for Strains; paired t test of AUC p = 0.0035). Meanwhile, administration of EcAZ-2^BSH, CmR^ either before or concurrently with EcAZ-2^SpecR^ led to equivalent co-colonization (**Fig. 3F,G**; Mixed Effects Model n.s.; paired t test of AUC n.s.). Thus, while AGD may have reduced the fitness of EcAZ-2^BSH, CmR^ when introduced after EcAZ-2^SpecR^, pre-engraftment or co-administration enabled both strains to occupy equivalent niche space. In conclusion, diet-induced changes in luminal metabolites can significantly impact the competition dynamics between isogenic strains.

### Inflammation-driven variation in LBT colonization dynamics

EcAZ-2^IL10, CmR^ was developed as a potential treatment of chronic gastrointestinal diseases, such as inflammatory bowel disease (IBD). However, IBD is a chronic relapsing disease, and thus it is unclear how timing of administration may impact colonization with the LBT. Using an acute dextran sodium sulfate (DSS)-induced colitis mouse model^37^, we gavaged three groups of mice with EcAZ-2^SpecR^ followed by a gavage of the IL10-expressing LBT a week later (**Fig. 4A**; **Fig. S3A-C**). In healthy control mice that did not receive DSS, EcAZ-2^IL10, CmR^ co-colonized with EcAZ-2^SpecR^ at 2 orders of magnitude lower levels than the parental strain (**Fig. 4B**; Mixed Effects Model p < 0.0001 for strains; paired t test of AUC p = 0.0001; **Fig. S3D**), replicating our previous findings (**Fig. 2D**). To determine the effects of LBT administration during a period of remission, mice were treated with EcAZ-2^IL10, CmR^ four days prior to DSS-colitis induction. In these mice, EcAZ-2^IL10, CmR^ initially colonized at lower levels than EcAZ-2^SpecR^ (**Fig. 4C**; Mixed Effects Model p < 0.0001 for strains; paired t test of AUC p = 0.0441). Colonization remained stable for approximately 10 days post-DSS cessation before rapidly declining to undetectable levels in the stool and GI tissue (**Fig. 4C**, **Fig. S3E**), indicating that a preventative administration of the LBT may not be suitable for long-term engraftment. To determine the effects of LBT administration during a flare up, mice were treated with EcAZ-2^IL10, CmR^ two days after colitis induction; EcAZ-2^IL10, CmR^ achieved and maintained equivalent co-colonization even after DSS cessation (**Fig. 4D**; Mixed Effects Model n.s.; paired t test of AUC n.s.; **Fig. S3F**). This suggests that inflammation during administration supports engraftment but post-engraftment inflammation can lead to LBT clearance.

**Figure 4.**
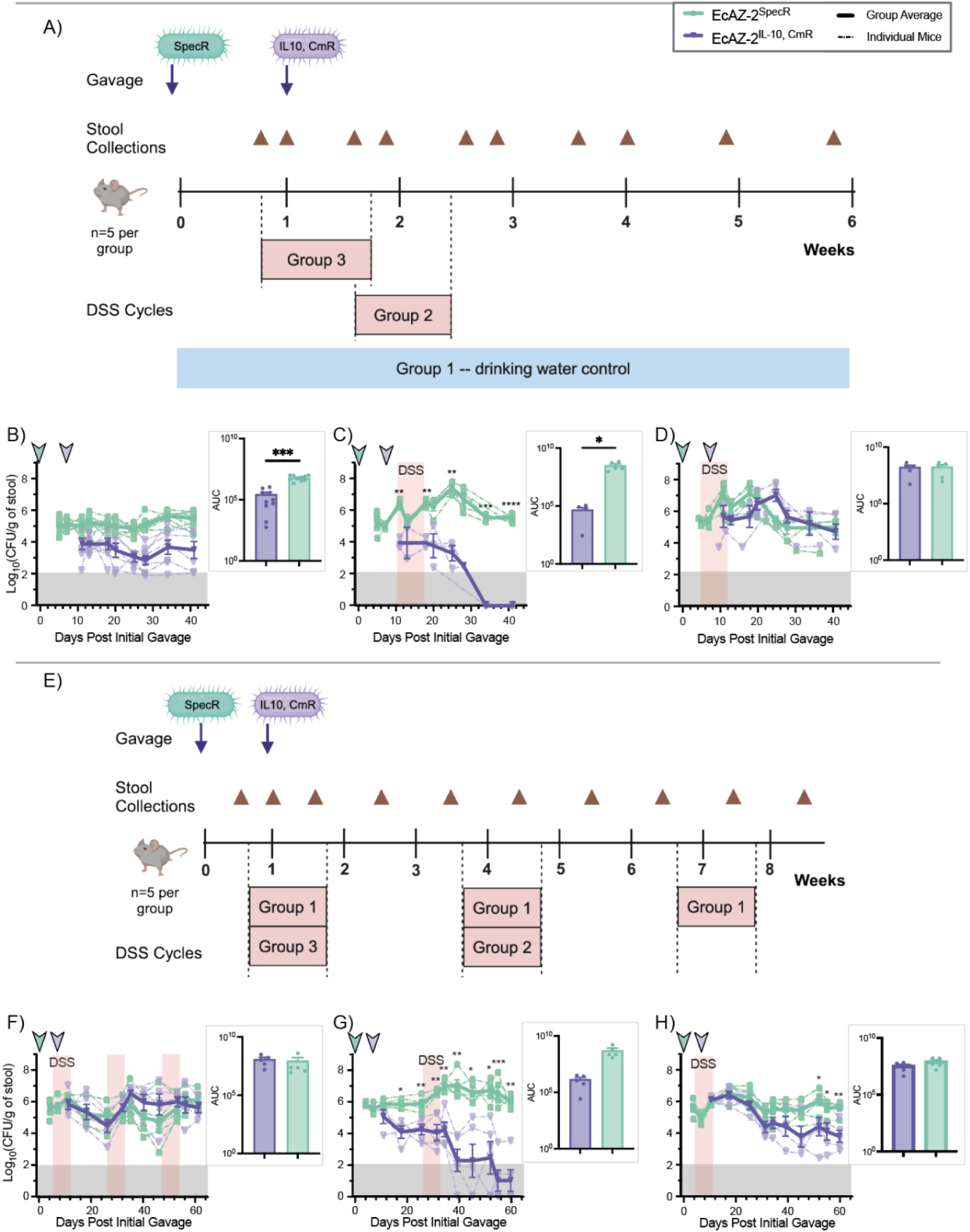
Inflammation-driven variation in LBT colonization dynamics. **(a)** Experimental design to test EcAZ-2^IL10, CmR^ competition against EcAZ-2^SpecR^ during acute DSS-induced colitis. Arrows indicate when the oral gavages occur. Brown triangles represent time points for stool collections. Red boxes with “Group #” represent when 2% DSS was administered in their drinking water. **(b-d)** Colonization curves for EcAZ-2^IL10, CmR^ (purple) and EcAZ-2^SpecR^ (green) competition in **(b)** control healthy mice, **(c)** when EcAZ-2^IL10, CmR^ is administered as a preventative, and **(d)** when EcAZ-2^IL10, CmR^ is administered as a treatment. **(e)** Experimental design to test EcAZ-2^IL10, CmR^ competition against EcAZ-2^SpecR^ during acute and chronic DSS-induced colitis. Arrows indicate when the oral gavages occur. Brown triangles represent time points for stool collections. Red boxes with “Group #” represent when 2% DSS was administered in their drinking water. **(f-h)** Colonization curves for EcAZ-2^IL10, CmR^ (purple) and EcAZ-2^SpecR^ (green) competition under **(f)** chronic relapsing colitis, **(g)** when EcAZ-2^IL10, CmR^ is administered as a preventative for acute colitis, and **(h)** when EcAZ-2^IL10, CmR^ is administered as a treatment for acute colitis. Arrowheads indicate when each strain was gavaged. Gray area indicates limit of detection. Translucent red shading indicates active DSS administration. AUC was calculated and compared across strains. Values are represented as mean ± the standard error of mean. Time points with asterisks along the colonization curves indicate significantly different levels in colonization between the two strains as determined by a Mixed Effects Model with a post-hoc Fisher’s LSD test. AUC comparisons were done via two-tailed paired t-test (* p < 0.05; ** p < 0.01; *** p<0.001; no asterisks = not significant).

To determine whether improved LBT colonization would be maintained upon additional flare ups, mice underwent three cycles of 1 week on 2% DSS followed by 2 weeks of normal drinking water (**Fig. 4E; Fig. S3G-I; Supplemental Table 2**). Encouragingly, EcAZ-2^IL10, CmR^ maintained equivalent co-colonization with EcAZ-2^SpecR^ upon subsequent cycles of colitis (**Fig. 4F**; Mixed Effects Model n.s.; paired t test of AUC n.s.). To determine if the previous clearance of the preventatively-administered LBT (**Fig. 1C**) was due to DSS disrupting engraftment, we repeated the preventative administration schema but first allowed EcAZ-2^IL10, CmR^ to establish colonization for 19 days before the onset of acute colitis. The results mimicked our initial findings (**Fig. 1C**; **Fig. 4G**; Mixed Effects Model p < 0.0001 for strains; paired t test of AUC n.s.), indicating that LBT clearance is due to shifts in the intestinal environment rather than initial engraftment dynamics. Administering EcAZ-2^IL10, CmR^ during acute colitis confirmed prolonged enhanced co-colonization with EcAZ-2^SpecR^ when tracked for a longer period of time (**Fig. 4H**; Mixed Effects Model n.s.; paired t test of AUC n.s.), before the colonization levels of EcAZ-2^IL10, CmR^ dropped to levels similar to that in healthy mice (**Fig. 1D, 4B**). In summary, intestinal inflammation at the time of EcAZ-2^IL10, CmR^ administration provides prolonged improved LBT colonization alongside a parental strain that is robust to subsequent inflammation challenges. Our results accentuate the adaptability of LBT engraftment while emphasizing local environmental conditions that need to be considered in future therapeutic applications^32^.

### Strategies for enhancing LBT colonization in competitive gut environments

After determining that EcAZ-2 LBTs consistently co-colonize with a parental strain in the gut, we explored strategies to increase LBT colonization levels for therapeutic efficacy. First, we tested increased LBT dosing by administering multiple gavages. Two groups of mice received EcAZ-2^SpecR^ followed by a gavage with EcAZ-2^BSH, CmR^ a week later. One group received 2 additional weekly gavages of EcAZ-2^BSH, CmR^ for a total of three doses. A single dose of EcAZ-2^BSH, CmR^ resulted in 1 order of magnitude lower colonization compared to EcAZ-2^SpecR^ (**Fig. 5A**; two-way ANOVA p = 0.0048 for strains; paired t test of AUC p = 0.0068). While three doses of EcAZ-2^BSH, CmR^ did not result in equivalent co-colonization with EcAZ-2^SpecR^ (**Fig. 5B**; Mixed Effects Model p = 0.0148 for strains; paired t test of AUC p = 0.0331), it did slightly but significantly increase overall LBT levels without altering levels of the parental strain (**Fig. S4A-B**; Mixed Effects Model, # of Admins p = 0.0273). We observed similar results with increased EcAZ-2^IL10, CmR^ doses. A single dose of EcAZ-2^IL10, CmR^ led to colonization about 3 orders of magnitude lower than that of EcAZ-2^SpecR^ (**Fig. 5C**; Mixed Effects Model p = 0.0221 for strains; paired t test of AUC p = 0.0051). While three doses of EcAZ-2^IL10, CmR^ did not result in equivalent co-colonization (**Fig. 5D**; Mixed Effects Model p = 0.057 for strains; paired t test of AUC p = 0.0068) it did increase EcAZ-2^IL10, CmR^ colonization levels by 1 order of magnitude (**Fig. S4C-D**; Mixed Effects Model, # of Admins p = 0.0122). Taken together, multiple doses significantly increased LBT colonization, even with an isogenic strain already engrafted.

**Figure 5.**
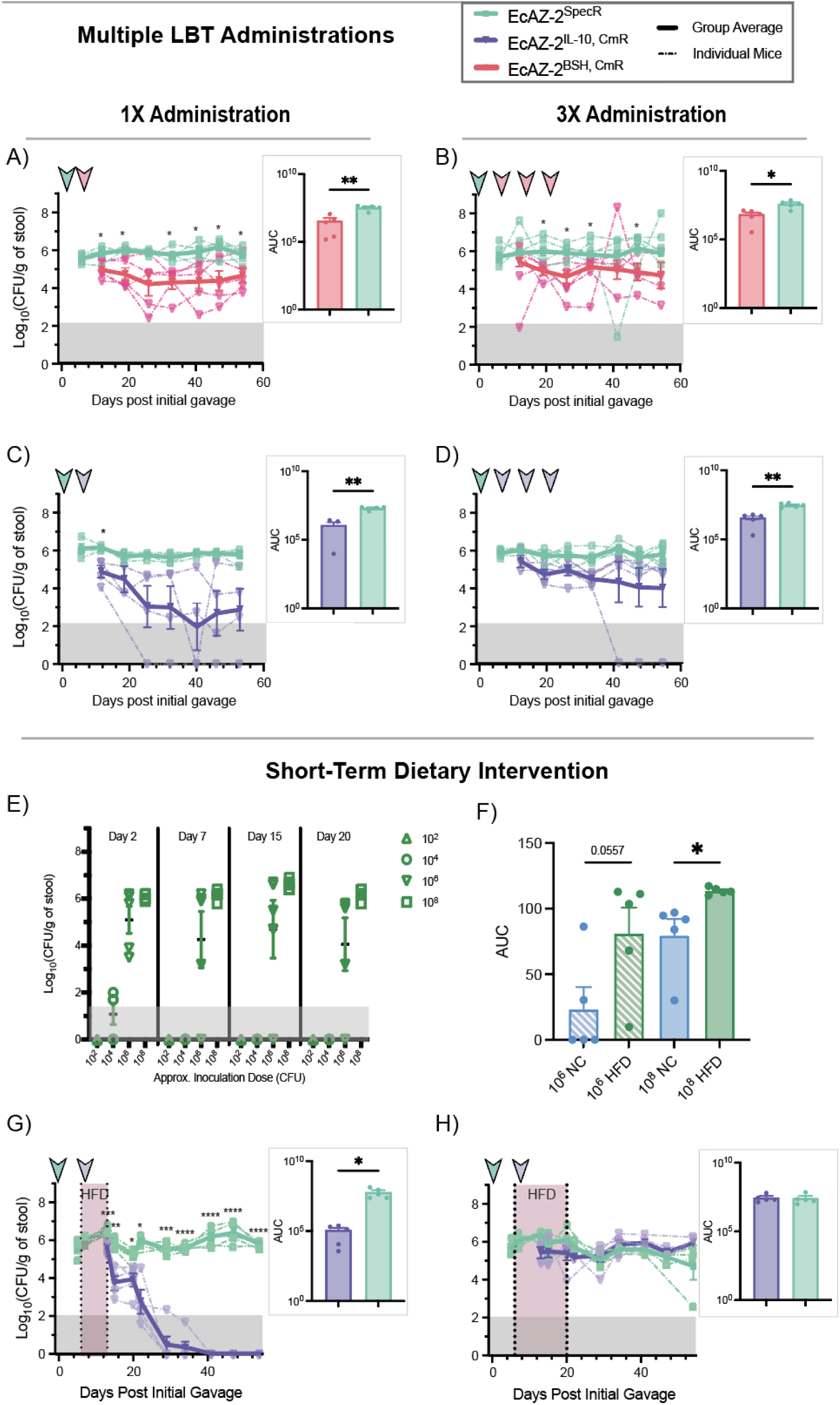
Strategies for enhancing LBT colonization in competitive gut environments. **(a-d)** Mice (n=4-5 per group) were first gavaged with EcAZ-2^SpecR^ followed by **(a-b)** EcAZ-2^BSH, CmR^ or **(c-d)** EcAZ-2^IL10, CmR^ one week later. **(b, d)** Two groups of mice (n=5) then received two additional gavages of their respective LBT for a total of three doses. Colonization curves for EcAZ-2^BSH, CmR^ (pink), EcAZ-2^IL10, CmR^ (purple), and EcAZ-2^SpecR^ (green) competition in SPF mice. Arrowheads indicate when each strain was gavaged. Gray area indicates limit of detection. AUC was calculated and compared across strains. **(e)** Four groups of mice (n = 5) were acclimated to HFD then received one gavage of 10^2^, 10^4^, 10^6^, or 10^8^ CFU of EcAZ-2. Colonization was tracked for 20 days to determine the minimum critical load for engraftment. **(f)** AUC was calculated for mouse groups receiving 10^6^ or 10^8^ CFU on HFD or normal chow (NC). NC data was reanalyzed from Russell *et al.* **(g-h)** Two groups of mice (n = 5 per group) were gavaged with EcAZ-2^SpecR^ (green), followed with EcAZ-2^IL10, CmR^ (purple) one week later and colonization was tracked. Starting one day before EcAZ-2^IL10, CmR^ administration, one group was switched to HFD for 1 week while the other group **(h)** was given HFD for 2 weeks. Arrowheads indicate when each strain was gavaged. Gray area indicates limit of detection. Translucent maroon shading indicates HFD administration. AUC was calculated and compared across strains. Values are represented as mean ± the standard error of mean. Time points with asterisks along the colonization curves indicate significantly different levels in colonization between the two strains as determined by a Mixed Effects Model with a post-hoc Fisher’s LSD test. AUC comparisons were done via two-tailed paired t-test (* p < 0.05; ** p < 0.01; *** p<0.001; no asterisks = not significant).

We next explored whether dietary intervention could modulate LBT colonization. Given that HFD increased the colonization levels of EcAZ-2-based strains (**Fig. 3B-D**), we first investigated its impact on the critical load of bacteria required for engraftment. Mice received either 10^2^, 10^4^, 10^6^, or 10^8^ CFU of EcAZ-2 and colonization was tracked for 20 days. Mice gavaged with 10^2^ or 10^4^ CFU had no detectable engraftment after day 2 (**Fig. 5E**). In contrast, 4 out of 5 mice receiving 10^6^ CFU and all mice receiving 10^8^ CFU maintained colonization (**Fig. 5E**). When reanalyzing previously gathered data for critical load in mice on normal chow^8^, we found that HFD significantly increased the AUC of EcAZ-2 in the 10^8^ CFU group (**Fig. 5F**; unpaired t test of AUC p = 0.0272). Additionally, the 10^6^ CFU group had increased AUC with HFD (**Fig. 5F**; unpaired t test of AUC p = 0.0557), though the results were not significant. Notably, 4 out of 5 mice on HFD for the same CFU group maintained colonization through Day 20 versus only 1 out of 5 on normal chow (**Fig. 5E**). This confirmed that HFD increases EcAZ-2 colonization and alters the critical load of bacteria required to colonize the gut.

We then employed a short-term HFD intervention one day before LBT gavage to try to boost EcAZ-2^IL10, CmR^ levels without allowing EcAZ-2^SpecR^ to outcompete it. Mice received EcAZ-2^SpecR^, followed by EcAZ-2^IL10, CmR^ a week later, and were kept on HFD for either 1 or 2 weeks before returning to normal chow. Surprisingly, 1 week of HFD was detrimental to LBT engraftment, ultimately leading to EcAZ-2^IL10, CmR^ clearance upon HFD discontinuation (**Fig. 5G**; Mixed Effects Model p < 0.0001 for strains; paired t test of AUC p = 0.0438). This mimicked patterns observed for EcAZ-2^IL10, CmR^ preventative administration before colitis onset (**Fig. 4C,G**). In contrast, 2 weeks of HFD led to sustained co-colonization of EcAZ-2^IL10, CmR^ and EcAZ-2^SpecR^, lasting even after mice returned to a normal chow diet (**Fig. 5H**; Mixed Effects Model n.s.; paired t test of AUC n.s.). Again, the competition dynamics mimicked those previously seen in colitis, but this time for the treatment administration of EcAZ-2^IL10, CmR^ (**Fig. 4D,H**). Together, this suggests that a threshold of inflammation must be achieved to support LBT engraftment^38–40;^ insufficient inflammation may trigger immune-mediated clearance of EcAZ-2^IL10, CmR^. Nonetheless, short-term dietary modifications can increase LBT colonization with a competitor strain already present.

## DISCUSSION

Our study challenges the long-standing assumption that the presence of one strain of a bacterial species invariably prevents engraftment of a secondary strain^14,17^. We demonstrated that EcAZ-2-based LBTs can co-colonize with a parental competitor in an intact microbiome (**Fig. 1G-I**; **Fig. 2E-G**). This was validated using both EcAZ-2^BSH, CmR^ and EcAZ-2^IL10, CmR^, highlighting the feasibility of using native bacterial chassis for effective LBT development and deployment. Importantly, competition outcomes in gnotobiotic mice (**Fig. 1D-F**) differed substantially from those in SPF mice, emphasizing the importance of studying microbiome-based therapeutics in intact, complex microbial communities. Our findings also revealed that strain arrival order and transgene-driven effects are key determinants of competitive success, emphasizing the need for preliminary testing to assess how therapeutic functions impact bacterial fitness and competition dynamics.

We identified several modifiable factors–host diet, dosing frequency, and intestinal inflammation–that can enhance LBT engraftment in competitive gut environments. While both HFD and AGD increase total bile acid concentration^33–36^ in the lumen, their effects on competition dynamics varied. HFD promoted increased colonization of EcAZ-2^SpecR^ (**Fig. 3B-3D**), while AGD enhanced the competitive advantage of the parental strain when it was introduced first (**Fig. 3E-G**). These findings demonstrate the complexity of diet-microbiome interactions and the challenges of predicting outcomes based on individual luminal metabolites. Guided by these insights, we implemented a two-week HFD intervention that significantly bolstered LBT colonization without promoting overgrowth of the parental strain (**Fig. 5H**). Similarly, increasing LBT dosing frequency modestly improved LBT colonization levels (**Supp fig**). These results suggest that engineered LBTs may need to be paired with a diet that optimizes their engraftment or efficacy.

One of the most striking findings was the role of active intestinal inflammation in shaping LBT colonization stability. Administering EcAZ-2^IL10, CmR^ during colitis led to equivalent co-colonization with the parental strain, with levels persisting after inflammation resolved (**Fig. 4D,H**). Conversely, administering the LBT prior to colitis onset often led to bacterial clearance upon resolution of inflammation (**Fig. 4C,G**). While the exact mechanisms remain unclear, we hypothesize that without active inflammation, the LBT strain can only occupy niches that are then destroyed when DSS-colitis is induced; this drastic change in colonization is likely to be host-mediated as DSS does not alter the microbiota in the absence of a host^41^. However, active inflammation creates new ecological niches which the engineered LBT can occupy equally well as the parental strain. Remarkably, subsequent DSS challenges did not disrupt EcAZ-2^IL10, CmR^ colonization when the LBT was administered during colitis (**Fig. 4F**). These findings suggest that IL10-expressing LBTs hold promise for long-term IBD treatment if administered during an active flare.

Collectively, our results demonstrate that competitive exclusion is not a fundamental barrier to LBT development using native bacterial chassis. By identifying and manipulating factors such as diet and dosing strategies, we provide actionable methods to optimize LBT colonization and therapeutic efficacy in complex gut environments.

Much of the foundational work in microbiome research has relied on simplified and controllable experimental models^17,24,42^. Testing LBTs *in vivo* often involves pre-treating SPF mice with antibiotics (e.g. streptomycin) to deplete the microbiome or using gnotobiotic mice colonized with defined microbial consortia. While these models offer experimental control, they fail to replicate the complexity of intact microbiomes, limiting their relevance to human applications. Mice with reduced communities lack key bacteria-to-bacteria and bacteria-to-host interactions, experience shifts in luminal metabolites, and exhibit changes in niche structure and availability^43^–all of which significantly impact LBT colonization dynamics^44–46^. These limitations are especially pronounced in germ-free mice, which have altered gastrointestinal morphology^47,48^ and underdeveloped immune systems^49,50^. Consistent with these deviations, we observed notable differences in competition of EcAZ-2^BSH, CmR^ against pre-established EcAZ-2^SpecR^ in gnotobiotic versus conventionally-raised mice (**Fig. 1D,G**). Thus, we advocate for prioritizing increasingly complex systems, including intact microbiomes, to enhance the translational relevance of microbiome research.

Previous studies of bacterial competition in the gut have largely relied on gnotobiotic mice. A seminal 2013 study of *Bacteroides* spp.^17^ concluded that two strains of the same species could not co-colonize in the gut of germ-free mice, while strains from different species could. However, in the context of more complex microbiomes, it is plausible that *Bacteroides* spp. strains might co-colonize, similar to the behavior of EcAZ-2 strains in our study. Indeed, recent studies using gnotobiotic mice colonized with complex synthetic microbiomes^24^ revealed coexistence for *Bacteroides vulgatus* in a ‘negative’ microbiome lacking additional *B. vulgatus* strains. In contrast, *Akkermansia municiphila* exhibited competitive exclusion under similar conditions^24^. These findings highlight the importance of microbiome complexity in shaping competition outcomes and underscore the need for further research in intact microbiomes.

Our work extends these insights by demonstrating that LBT colonization is shaped by multiple host and environmental factors, including diet, inflammation, and dosing. While this study focused on a single strain of *E. coli*, which makes up less than 0.1% of the murine gut microbiome^8^, future research should explore diverse bacterial species and strains to evaluate how these factors influence colonization dynamics across the microbiome. Furthermore, it should be considered that the gut microbiome composition of laboratory-raised mice differs from that of free, wild mice^51^–these variations should also be explored.

Additionally, while this study examined LBTs constitutively-expressing single-gene therapeutics, future work should investigate the impact of inducible function expression and more complex genetic circuits^52–56^ to broaden the therapeutic potential of LBTs.

Finally, as microbiome research advances, the development of “smarter” LBTs represents a promising frontier^57^. These LBTs would be designed to integrate into the gut microenvironment, leveraging identified host and environmental factors to enhance engraftment^58–61^. Engrafted LBTs offer unique advantages, including long-term localized delivery of therapeutic compounds, enabling the use of labile drugs that would otherwise degrade in the gastrointestinal tract. Local delivery also reduces the required therapeutic dose, minimizing systemic side effects^62^. Ultimately, the most transformative LBTs could be engineered to sense disease biomarkers and respond dynamically, producing therapeutic compounds proportional to biomarker levels^57^. This precision medicine approach has the potential to revolutionize treatment for chronic and complex diseases, offering durable solutions that integrate seamlessly into the host microbiome.

## METHODS

### RESOURCE AVAILABILITY

#### Lead contact

Further information and requests for resources should be directed to and will be fulfilled by the lead contact, Amir Zarrinpar.

#### Materials availability

EcAZ-2, EcAZ-2^BSH, CmR^, EcAZ-2^IL10, CmR^, and EcAZ-2^SpecR^ will be made available subject to a materials transfer agreement with the University of California, San Diego.

#### Code and Data Availability

This paper does not report original code. Genome sequencing will be deposited in ENA (project accession number PRJEB85101 and data used to generate figures is being deposited on Figshare. Any additional information required to reanalyze the data reported in this paper is available from the lead contact upon request.

### EXPERIMENTAL MODEL DETAILS

#### Bacterial strains

For the construction of EcAZ-2^SpecR^, the spectinomycin resistance gene *aadA* was inserted between genes *murB* (UDP-N-acetylenolpyruvoylglucosamine reductase) and *murI* (glutamate racemase) via lambda red recombineering (pSIM18). Spectinomycin resistant transformants were then used to inoculate luria broth (LB) cultures containing selective antibiotics. Genomic DNA was isolated using the Zymo Quick-DNA Miniprep Plus Kit and concentrated to 50ng/uL using AMPure beads, according to manufacturer instructions. Genomic DNA was sent to Plasmidsaurus for Nanopore long-read sequencing and subsequent assembly. The assembled genomes were then compared to the index EcAZ-2 genome to confirm successful insertion of *aadA* without off-target effects. All EcAZ-2 based strains were cultured in Luria Broth or Super Optimal Broth at 37C with shaking.

#### J774A.1 Cells

J774A.1 macrophage-like cells, isolated from the ascites of a female mouse with reticulum cell sarcoma, were cultured at 37C with 5% CO2 in Dulbecco’s Modified Eagle Medium (DMEM) supplemented with 10% sterile fetal bovine serum. These cells were used in the IL-10 bioactivity assay detailed below.

#### Gnotobiotic Mice

Germ-free C57BL/6 mice were born and reared in flexible film isolators and maintained under gnotobiotic conditions at the University of Nebraska-Lincoln. All mice were fed an autoclaved chow diet (LabDiet 5K67, Purina Foods, St. Louis, MO) ad libitum. The Institutional Animal Care and Use Committee (IACUC) at the University of Nebraska-Lincoln approved all procedures involving animals (protocol 2126).

After being transferred to a Tecniplast IsoCage unit, 15 male C57BL/6 mice, around 10 weeks of age, were inoculated with ∼10^10^ colony forming units (CFU) of bacteria resuspended in 200uL of 1x PBS via oral gavage on day 0. The mice were split into 3 groups: group 1 received EcAZ-2^SpecR^, group 2 received EcAZ-2^BSH^, and group 3 received a 50/50 mixture of EcAZ-2^SpecR^ and EcAZ-2^BSH^. One week later, all mice received a second gavage of ∼10^10^ CFU of bacteria in 200uL of PBS. Group 1 received EcAZ-2^BSH^, group 2 received EcAZ-2^SpecR^, and group 3 received another 50/50 mixture of EcAZ-2^SpecR^ and EcAZ-2^BSH^.

Colonization was monitored by collecting and weighing ∼3 fecal pellets per mouse at day 10, 15, 28, and 49. Stool was resuspended in 1mL of deionized (DI) water and further diluted to 10^-7^ in DI water. 100uL was plated on LB agar plates containing selective antibiotics for either EcAZ-2^SpecR^ (spectinomycin 100ug/mL) or EcAZ-2^BSH^ (chloramphenicol 40ug/mL) and plates were incubated at 37C overnight. The next day colonies were counted and the CFU per gram of stool for each strain was calculated.

#### Specific-Pathogen Free Mice

All animal experiments were conducted in accordance with the guidelines of the IACUC of the University of California, San Diego. C57BL/6 mice (Jackson Laboratories) were housed in a specific pathogen free facility (irradiated chow and autoclaved bedding). Mice were housed 3-5 per cage unless otherwise stated. All mice were fed a normal-chow diet (NC; Diet 7912, Teklad Diets, Madison, WI), unless specifically indicated to be on high fat diet (HFD; Diet D12492i, Research Diets, New Brunswick, NJ) or atherogenic diet (AGD; Diet TD.96121, Teklad Diets, Madison, WI). For mice on a diet other than NC, a 2 week dietary acclimation period occurred before bacterial gavage.

5-8 week old male C57BL/6 mice were inoculated with ∼10^10^ CFU of bacteria resuspended in 200uL of 1x PBS via oral gavage on day 0. Typically, 15 mice per experiment were split into three groups: group 1 received bacterial strain 1, group 2 received bacterial strain 2, and group 3 received a 50/50 mixture of strain 1 and 2. One week later, each group received a second bacterial gavage: group 1 received bacterial strain 2, group 2 received bacterial strain 1, and group 3 received another 50/50 mixture of strain 1 and 2. Colonization levels of the two strains were monitored by stool collection on roughly a weekly basis (detailed below). EcAZ-2^SpecR^ colonization was measured on LB agar plates containing spectinomycin. EcAZ-2^BSH^ or EcAZ-2^IL10^ colonization was measured on LB agar plates containing chloramphenicol.

### METHOD DETAILS

#### Colonization Assessment

For stool colonization, we collected 3-5 stool pellets from individual mice and weighed them. 1 mL of sterile deionized water was added to each sample along with one sterile chrome bead (Neta Scientific, Hainesport, NJ). Samples were homogenized for 20 seconds. at 4500 rpm in Mini-Beadbeater-24 (Biospec, Bartlesville, OK). Samples were diluted in DI water and plated on chloramphenicol plates and spectinomycin plates to calculate CFU/gram of stool.

For tissue colonization, upon sacrifice, the intestines were removed and separated into the duodenum, jejunum, terminal ileum, cecum, and transverse colon. Luminal contents were roughly removed by squeezing, with no intestinal flushing. A small portion of each tissue section was sampled, weighed, and homogenized in 1mL of sterile DI water using 2 chrome beads and two rounds of bead beating as detailed above. Appropriate dilutions of the homogenized tissues were then plated on selective agar plates to calculate CFU/gram of tissue for each strain at each GI tract location.

#### Cerillo Duet In Vitro Competition

Indirect competition between bacterial strains was conducted using the Cerillo Duet system. Each strain was grown in triplicate in rich SOB with appropriate antibiotics overnight. The next morning, overnight broth cultures were used to prepare a 1:10 dilution in SOB broth with only kanamycin (12.5ug/mL). 800uL of SOB broth + kanamycin was added to either side of each membrane and 8uL of the appropriate 1:10 diluted strain was inoculated on either side; each pairwise growth comparison was inoculated in triplicate. An 18 hour kinetic growth curve was measured via OD600 on a Tecan; the plate was incubated at 37C and measurements were taken every 10 minutes after 5 seconds of linear shaking, amplitude 3mm.

For Duet competition in the presence of bile acids, overnight broth cultures of each strain were grown in SOB with appropriate antibiotics in triplicate. The next morning. 800uL of SOB with 0.15% bile salts was added to either side of each Duet. 4uL of overnight cultures were used to inoculate either side of the appropriate Duets; each pairwise growth comparison was inoculated in triplicate. An 18 hour kinetic growth curve was measured via OD600 on a Tecan; the plate was incubated at 37C and measurements were taken every 10 minutes after 5 seconds of linear shaking, amplitude 3mm.

#### DSS-Induced Chemical Colitis in Mice

2% of dextran sodium sulfate (DSS) was administered via drinking water for 7 days to the SPF C57BL/6 mice to induce acute colitis. “Treatment” administration of EcAZ-2^IL10^ was conducted by oral gavage of ∼10^10^ CFU in 0.2mL of PBS 2 days after starting DSS administration. “Preventative” administration of EcAZ-2^IL10^ was conducted by oral gavage either 4 or 19 days before DSS administration.

Chronic colitis was mimicked using 3 rounds of 7 days of 2% DSS, followed by 2 weeks of normal water between DSS rounds. For chronic colitis, EcAZ-2^IL10^ was administered via oral gavage 2 days after the start of the first round of DSS, aligned with the acute colitis “treatment” administration paradigm.

Body weights were monitored daily for starting the day before DSS administration began and continuing for a total of 9 days to monitor for >20% drop in body weight. If any mice lost more than 20% of their body weight, they would be euthanized. Stool collections were conducted at least once a week to track colonization levels of each strain.

#### Critical Inoculation Dose Determination

Mice were acclimated to HFDt for 2 weeks prior to bacterial gavage. All mice received a 0.2mL oral gavage in PBS of approximately 10^2^, 10^4^, 10^6^, or 10^8^ CFU of native *E. coli* chassis EcAZ-2. Engraftment levels were monitored by stool collection as detailed above and plated on LB agar plates containing kanamycin at 2, 7, 15, and 20 days after oral gavage. Comparison to normal chow critical inoculation results was conducted via reanalyzing data previously published in Russell *et al*. *Cell* 2022.

#### Short-Term Dietary Intervention for Engraftment

Mice were maintained on normal chow diet, outside of the defined dietary intervention period. Mice were gavaged with 10^10^ CFU of EcAZ-2^SpecR^ at day 0. Six days after the first gavage, mice receiving the dietary intervention were switched to HFD. One day later (seven days post the first gavage), mice received a gavage of 10^10^ CFU of EcAZ-2^IL10^. One group of mice that was switched back to NC after 1 week and the other after 2 weeks of HFD. Stool collections were conducted weekly to track colonization levels of each strain.

#### Multiple Administration for Engraftment

Mice received a gavage of 10^10^ CFU of the parental EcAZ-2^SpecR^ chassis in 0.2mL PBS at day 0. 1 week later, all mice received a gavage of either the EcAZ-2^BSH^ or EcAZ-2^IL10^ LBT. Half of the mice receive 2 more subsequent gavages of the same LBT as before, all spaced one week apart, for a total of 1 or 3 administrations of each LBT. Stool collections were conducted weekly to track colonization levels of each strain.

#### BSH Functional Maintenance Assessment

Functional maintenance of the BSH therapeutic product after EcAZ-2^BSH^ engraftment in the murine gut was conducted using LB agar plates containing 40ug/mL of chloramphenicol and 8.6mM taurine deoxycholic acid (TDCA). BSH deconjugation activity was determined qualitatively by observing precipitate formation surrounding a colony that results from the deconjugation of taurine from TDCA.

#### IL10 Functional Maintenance Assessment

Functional maintenance of the IL10 therapeutic product after EcAZ-2^IL10^ engraftment was determined by measuring the presence of IL10 bioactivity. Glycerol stocks of homogenized stool from the final stool collection of EcAZ-2^IL10^ competition with EcAZ-2^SpecR^ were used to inoculate 2mL of SOB + 40ug/mL chloramphenicol. Cultures were incubated at 37C with shaking overnight. The next morning, OD600 of each culture was measured and the amount of overnight culture needed to create 700uL of PBS with an OD600=1.0 was determined and spun down at 4000xg for 10 minutes in a 1.7mL tube. The resultant pellet was resuspended in 700uL of PBS and lysed via sonication (Fisher Scientific Model 120 Sonic Dismembrator) using 10 cycles of lysis at 50% amplitude for 20 seconds on followed by 20 seconds off. Lysate was aliquoted and stored at -70C until later analysis.

The amount of IL10 protein in each lysate was determined using the Human IL10 ELISA Kit (Biolegend, Catalog #430604), following manufacturer instructions. The total amount of protein was determined using the BCA Protein Quantification Assay (Lamda Biotech Inc., Catalog #G1002) following the Microplate Protocol provided by the manufacturer.

IL10 bioactivity was determined using a macrophage cell assay. Briefly, 500uL of 1×10^5^ cells per mL of mouse macrophage-like cells J774.1 were seeded in a 24-well tissue culture plate and incubated for 2 hours at 37C in 5% CO_2_. 10uL of bacterial lysate for each mouse was added with 10uL of PBS to the appropriate wells. 10uL of EcAZ-2 lysate + 10uL of PBS served as the negative control. Positive controls included treatment with: 10uL of stock strain EcAZ-2^IL10^ + 10uL PBS, 10uL of EcAZ-2 lysate + 10uL of 100ng/mL recombinant IL10 (R&D Systems, Catalog #1064-ILB--010) in PBS,10uL of EcAZ-2 lysate + 10uL of 1ng/mL recombinant IL10 in PBS, and 10uL of EcAZ-2 lysate + 10uL of 0.5ng/mL recombinant IL10 in PBS. Samples and standards were incubated with cells for 4 hours at 37C with 5% CO_2_. Media was harvested and stored at -70C for further analysis. TNF-alpha production by the treated J774.1 cells was measured using harvested supernatants diluted 1:5 in PBS using the mouse TNF-alpha ELISA kit (Biolegend, Catalog #430904), following manufacturer instructions. EcAZ-2^IL10^ functional bioactivity was assessed qualitatively by determining if TNF-alpha production repression is greater than or equal to the repression by the stock strain of EcAZ-2^IL10^.

### QUANTIFICATION AND STATISTICAL ANALYSIS

All comparisons of colonization levels of EcAZ-2^BSH, CmR^ or EcA-2^IL10, CmR^ and EcAZ-2^SpecR^ were conducted using two methods. First, a Mixed Effects Model was run (due to missing values in the data) with a post-hoc Fisher’s LSD test to analyze log_10_(CFU/gram of stool) colonization levels of competing strains across time points; the first one (**Fig. 1G,H; 2E,F; 3C,D; 3F,G; 5B-D**) or first two (**Fig. 4B-D; 4F-H; Fig 5G,H**) stool collections were excluded if the second strain had yet to be administered. In a few cases where there were no missing values, a two-way ANOVA was used instead, due to its increased statistical sensitivity (**Fig. 1D-F, 2G, 3D, 5A**). Additionally, the AUC of CFU/gram of stool curves was calculated, omitting the first 2 or 3 stool collections respectively to remove artificially high ‘colonization’ data from strains that were recently orally administered and had yet to finish engrafting. A two-tailed paired t-test was run to compare the AUC of the LBT against the AUC of the parental strain. Likewise, AUC for the optical density of bacterial growth was calculated for each control and experimental Duet. A one-way ANOVA was run and each AUC was compared to the strain’s control AUC using Sidak multiple comparison corrections. A p-value < 0.05 was considered statistically significant. Finally, the AUC for the 10^6^ and 10^8^ CFU groups of mice that were either on normal chow diet (reanalyzed from Russell et al 2022 Cell) or HFD was calculated. Two-tailed t tests were run across diets. All graphs were made and statistical analysis was performed using GraphPad Prism software (version 10.4.0).

## ACKNOWLEDGEMENT

N.S. was supported by NSF GRFP and was a 2021 Biolegend Fellow. M.T. was partially supported by the UCSD Eureka! Scholars Award. A.Z. is supported by the CCF Litwin IBD Pioneers Program Award 1049147, VA Merit BLR&D Award I01 BX005707, and NIH R01 HL148801, R01 EB030134, R01 AI163483, and U01 CA265719. All authors receive institutional support from NIH P30 DK120515, P30 DK063491, P30 CA014195, and UL1 TR001442. The funders had no role in study design, data collection and interpretation, or the decision to submit the work for publication. The contents do not represent the views of the U.S. Department of Veterans Affairs or the United States Government.

## AUTHOR CONTRIBUTIONS

Conceptualization, N.S., M.T., A.Z.; Methodology, N.S., S.B., J.P., A.R-T., and A.Z.; Formal Analysis, N.S. and S.B.; Investigation, N.S., M.S., H.G., and S.B.; Resources, J.P., A.R-T., and A.Z.; Writing (Original Draft), N.S., M.S., and H.G.; Writing–Reviewing & Editing, N.S., S.B., M.S., H.G., M.T., I.M., J.P., A.R-T., and A.Z.; Visualization, N.S. and S.B.; Supervision, A.Z.; Project Administration, A.Z.; Funding Acquisition, A.Z.

## DECLARATION OF GENERATIVE AI AND AI-ASSISTED TECHNOLOGIES IN THE WRITING PROCESS

During the preparation of the work the authors N.S. and A.Z. used ChatGPT-4o to improve the clarity and readability of this manuscript. After using this tool, all authors reviewed and edited the content as needed and take full responsibility for the content of the published article.

