## Supplemental Figures and Tables for "Gut Competition Dynamics of Live Bacterial Therapeutics Are Shaped by Microbiome Complexity, Diet, and Therapeutic Transgenes"

********SUPPLEMENTARY MATERIAL********

Figure S1. EcAZ-2$^{BSH,\ CmR}$ shows no inherent fitness deficits compared to EcAZ-2$^{Spec}$
Figure S2. EcAZ-2$^{IL10,\ CmR}$ has slightly reduced fitness compared to EcAZ-2$^{SpecR}$
Figure S3. DSS-induced colitis differentially affects EcAZ-2$^{IL10,\ CmR}$ competition based on timing of LBT administration
Figure S4. Multiple administrations of EcAZ-2 LBTs increases LBT colonization levels
Table S1. BSH activity is maintained throughout EcAZ-2$^{BSH,\ CmR}$ competition with EcAZ-2$^{SpecR}$
Table S2. Colitis scoring of colons for mice at time of sacrifice

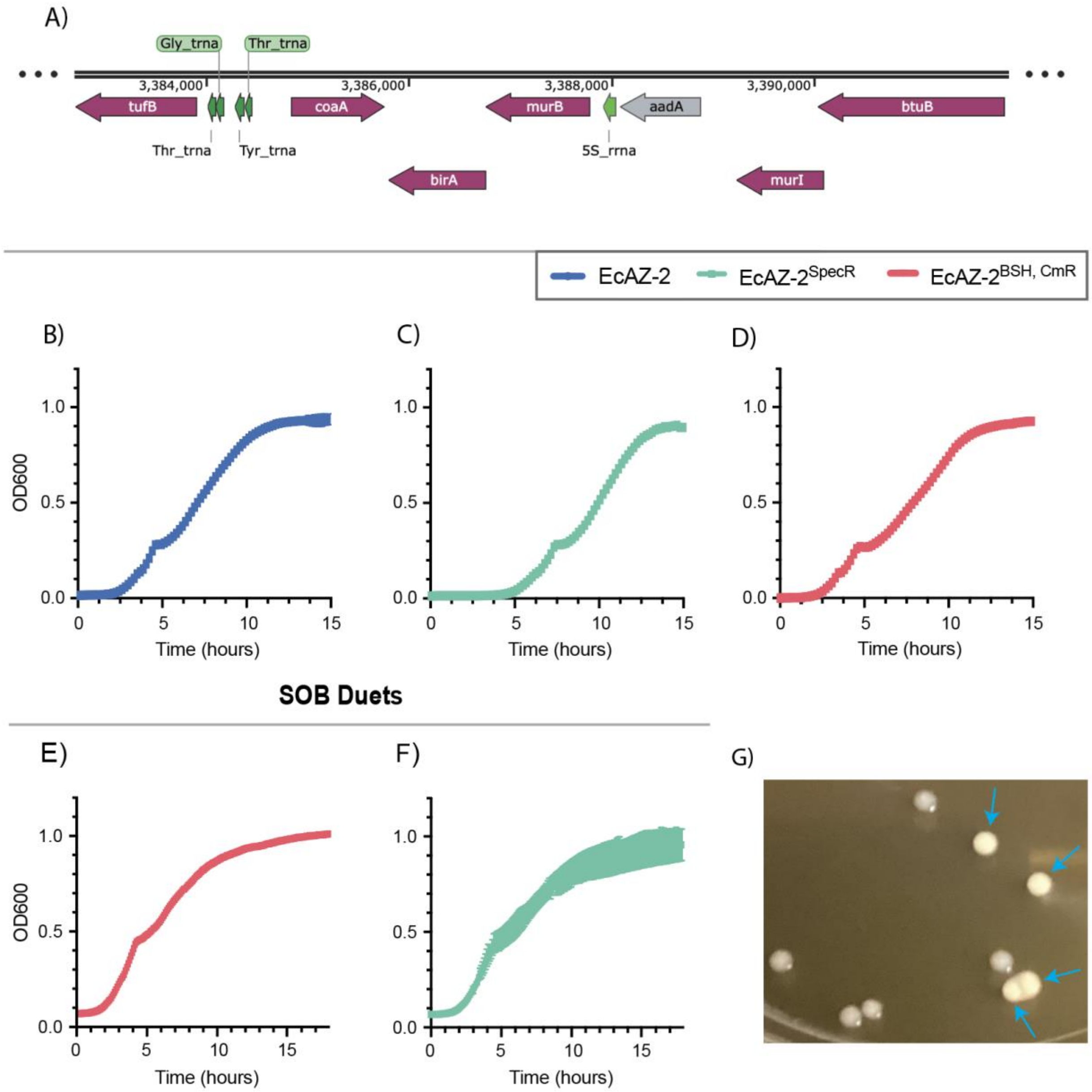


**Figure S1. EcAZ-2$^{BSH, CmR}$ shows no inherent fitness deficits compared to EcAZ-2$^{Spec}$, related to Figure 1 and Supplemental Table 1.** (**A**) Transgene insertion location of *aadA* (in gray) in EcAZ-2$^{SpecR}$ as determined by minION sequencing. Growth curves for (**B**) EcAZ-2, (**C**) EcAZ-2$^{SpecR}$, and (**D**) EcAZ-2$^{BSH, CmR}$. Duet growth of (**E**) EcAZ-2$^{BSH, CmR}$ and (**F**) EcAZ-2$^{SpecR}$ in SOB when growing against self. (**G**) Representative image of EcAZ-2 with and without BSH activity on an LB + TDCA plate. Blue arrows indicate colonies with BSH activity.

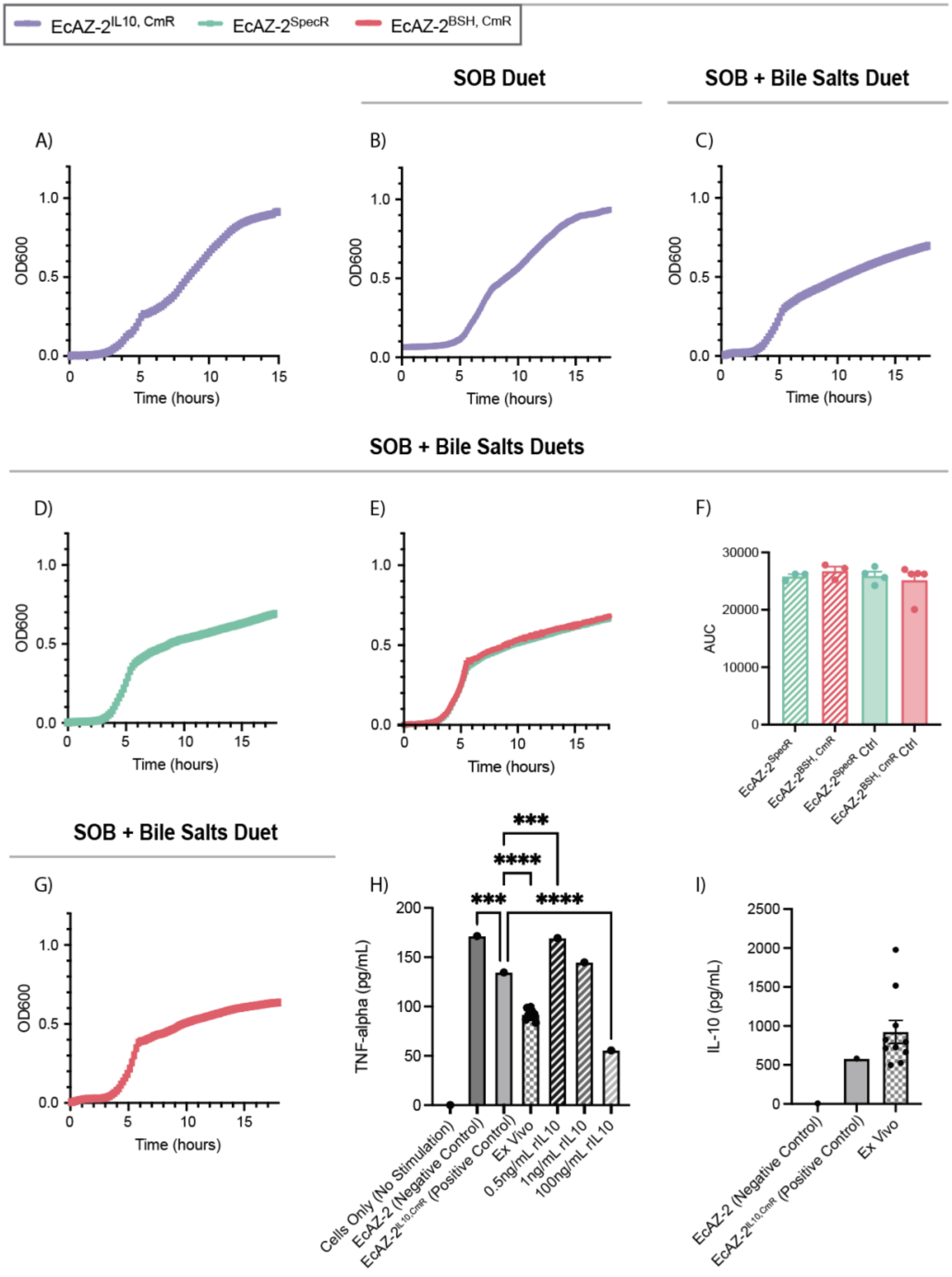


**Figure S2. EcAZ-2$^{\text{IL10, CmR}}$ has slightly reduced fitness compared to EcAZ-2$^{\text{SpecR}}$, related to Figure 2.** (**a**) Growth curve of EcAZ-2$^{\text{IL10, CmR}}$. (**b**) Duet growth of EcAZ-2$^{\text{IL10, CmR}}$ in SOB when growing against self. Duet growth of (**c**) EcAZ-2$^{\text{IL10, CmR}}$ and (**d**) EcAZ-2$^{\text{SpecR}}$ in SOB with bile salts when growing against self. (**e**) Duet growth of EcAZ-2$^{\text{BSH, CmR}}$ in SOB with bile salts when growing against EcAZ-2$^{\text{SpecR}}$ and (**f**) AUC of Duet optical density. (**f**) Duet growth of EcAZ-2$^{\text{BSH, CmR}}$ in SOB with bile salts when growing against self. (**g**) TNF-alpha production by stimulated mouse macrophage cell line J774A.1, reduced by IL10 anti-inflammatory bioactivity. (**h**) IL100=p-roduction per million cells after reisolation of a pooled LBT population from mice.

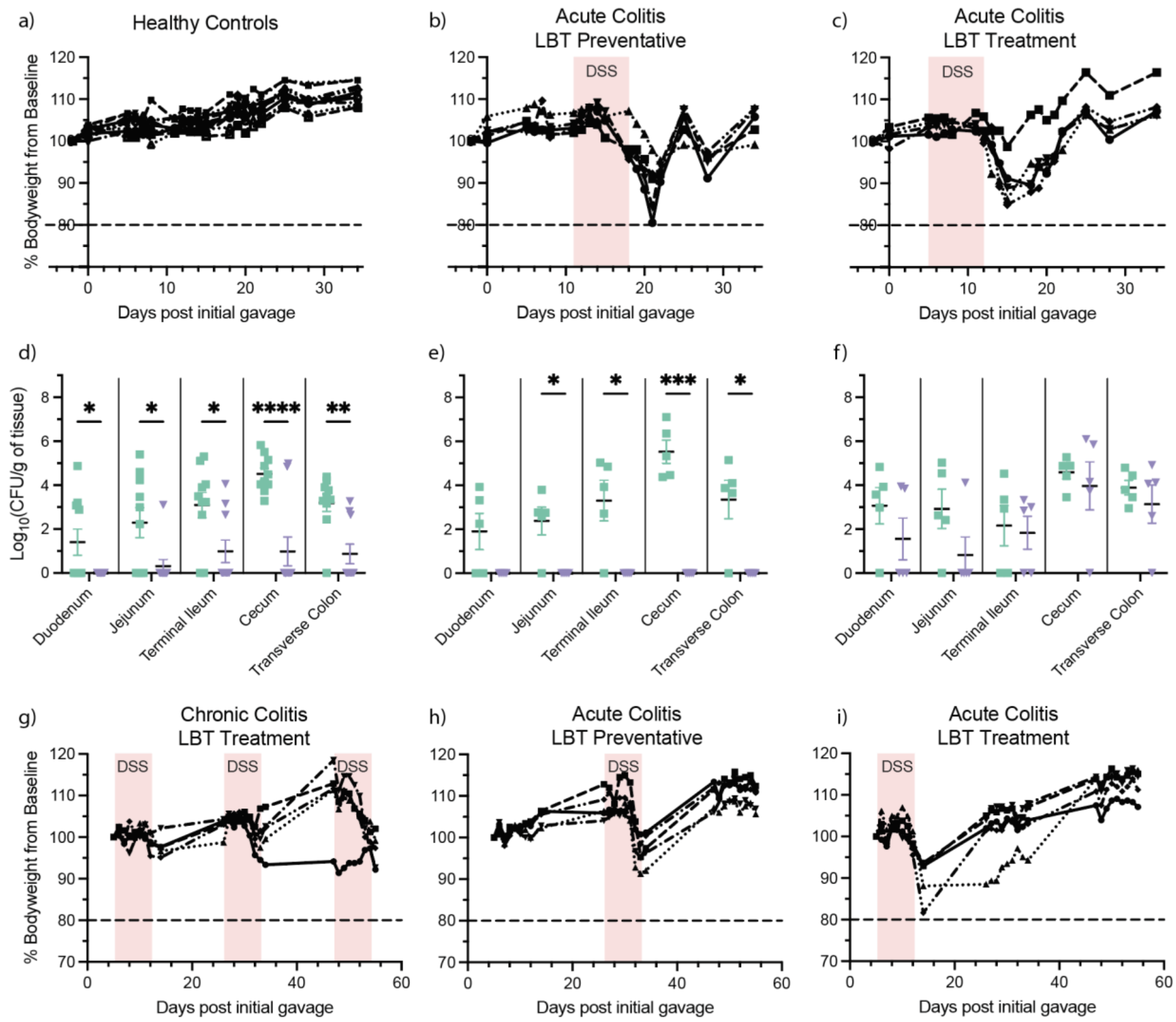


**Figure S3. DSS-induced colitis differentially affects EcAZ-2$^{IL10,\ CmR}$ competition based on timing of LBT administration, related to Figure 4 and Supplemental Table 2.** Percentage weight as compared to baseline for (**a**) healthy control mice, (**b**) mice undergoing acute colitis and receiving EcAZ-2$^{IL10,\ CmR}$ 4 days before colitis onset as a preventative, (**c**) and mice undergoing acute colitis and receiving EcAZ-2$^{IL10,\ CmR}$ 2 days after colitis onset as a treatment. Tissue colonization levels of EcAZ-2$^{SpecR}$ and EcAZ-2$^{IL10,\ CmR}$ in (**d**) healthy control mice, (**e**) mice receiving EcAZ-2$^{IL10,\ CmR}$ 4 days before colitis onset as a preventative, and (**f**) mice receiving EcAZ-2$^{IL10,\ CmR}$ 2 days after colitis onset as a treatment. Percentage weight as compared to baseline for (**g**) mice subjected to chronic colitis, (**h**) mice undergoing acute colitis and receiving EcAZ-2$^{IL10,\ CmR}$ 19 days before colitis onset as a preventative, (**i**) and mice undergoing acute colitis and receiving EcAZ-2$^{IL10,\ CmR}$ 2 days after colitis onset as a treatment.

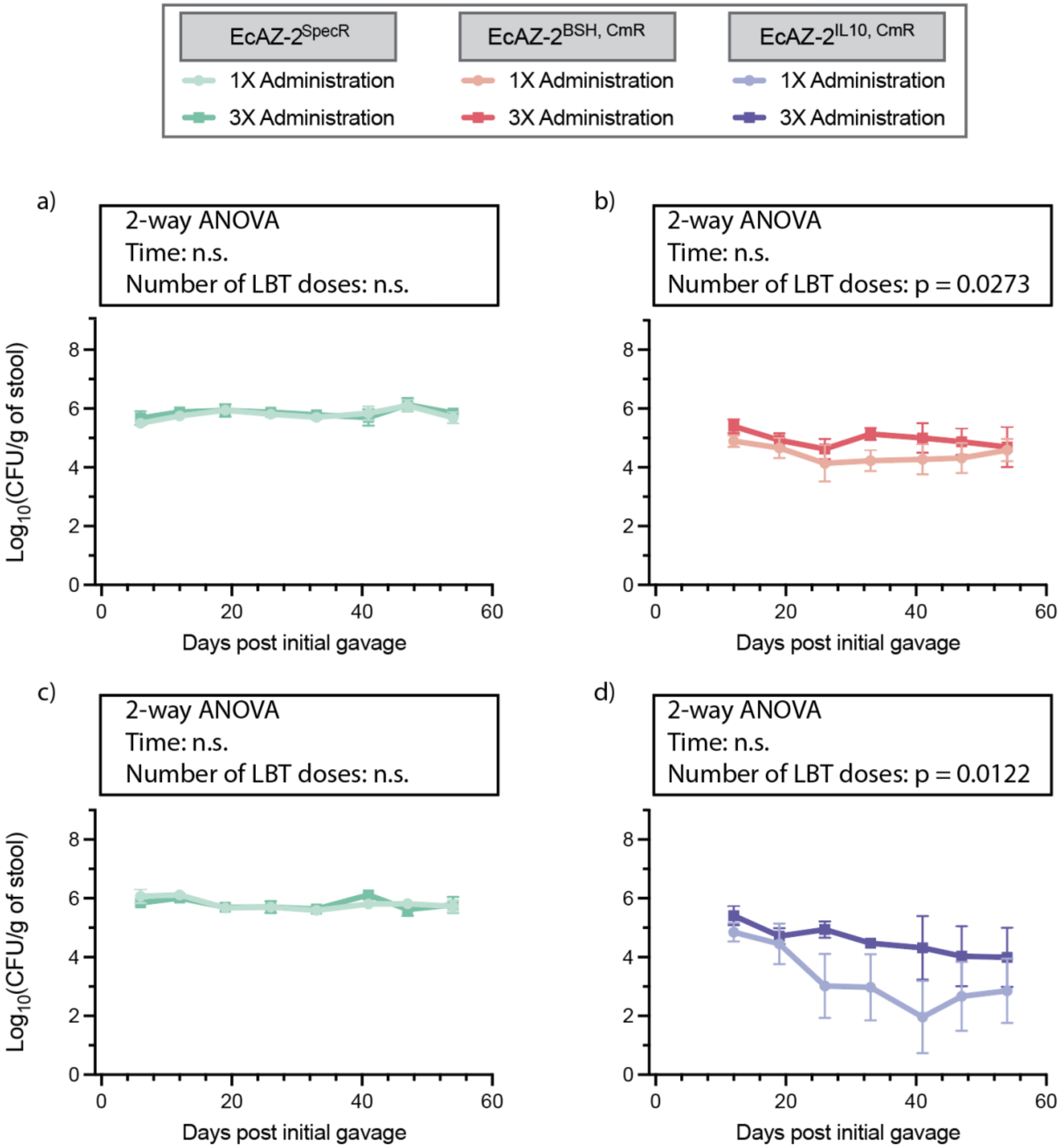


**Figure S4. Multiple administrations of EcAZ-2 LBTs increases LBT colonization levels, related to Figure 5.** Comparison of colonization levels of (**a**) EcAZ-2$^{SpecR}$ and (**b**) EcAZ-2$^{BSH, CmR}$ in mice receiving one versus three doses of the LBT. Comparison of colonization levels of (**c**) EcAZ-2$^{SpecR}$ and (**d**) EcAZ-2$^{IL10, CmR}$ in mice receiving one versus three doses of the LBT. 2-way ANOVA was run and p-values are listed on the graphs.

**Supplemental Table 1. BSH activity is maintained throughout EcAZ-2$^{BSH, CmR}$ competition with EcAZ-2$^{SpecR}$, as evaluated at the end of the study**

| Mouse ID | First Bacteria Received | Chloramphenicol Resistant Bacteria Detected? | BSH Deconjugation Activity Detected? |
|---|---|---|---|
| 1N | EcAZ-2$^{SpecR}$ | N | N |
| 1L | EcAZ-2$^{SpecR}$ | Y | Y |
| 1LL | EcAZ-2$^{SpecR}$ | N | Y |
| 1R | EcAZ-2$^{SpecR}$ | N | N |
| 1B | EcAZ-2$^{SpecR}$ | Y | Y |
| 2N | EcAZ-2$^{BSH, CmR}$ | Y | Y |
| 2LL | EcAZ-2$^{BSH, CmR}$ | Y | Y |
| 2R | EcAZ-2$^{BSH, CmR}$ | Y | Y |
| 2B | EcAZ-2$^{BSH, CmR}$ | Y | Y |
| 3N | EcAZ-2$^{SpecR}$ and EcAZ-2$^{BSH, CmR}$ | Y | Y |
| 3L | EcAZ-2$^{SpecR}$ and EcAZ-2$^{BSH, CmR}$ | Y | Y |
| 3LL | EcAZ-2$^{SpecR}$ and EcAZ-2$^{BSH, CmR}$ | Y | Y |
| 3R | EcAZ-2$^{SpecR}$ and EcAZ-2$^{BSH, CmR}$ | Y | Y |
| 3B | EcAZ-2$^{SpecR}$ and EcAZ-2$^{BSH, CmR}$ | Y | Y |

**Supplemental Table 2. Colitis scoring of colons for mice at time of sacrifice, related to Figure 4A-D**

| Mouse ID | DSS Administration | Inflammation | Crypt Abscesses | Inflammation Extent | Crypt Damage | Edema | Goblet Loss | Epithelial Hyperplasia | Total Score |
|---|---|---|---|---|---|---|---|---|---|
| 1N | Chronic | 2 | 0 | 1 | 1 | 1 | 0 | 1 | 6 |
| 1L | Chronic | 2 | 0 | 1 | 0 | 1 | 0 | 1 | 5 |
| 1R | Chronic | 2 | 0 | 1 | 1 | 1 | 0 | 1 | 6 |
| 1B | Chronic | 2 | 0 | 2 | 1 | 1 | 0 | 1 | 7 |
| 1LL | Chronic | 1 | 0 | 1 | 0 | 0 | 0 | 0 | 2 |
| 2N | Acute (LBT Treatment) | 2 | 0 | 1 | 0 | 1 | 0 | 1 | 5 |
| 2L | Acute (LBT Treatment) | 1 | 0 | 1 | 0 | 0 | 0 | 0 | 2 |
| 2R | Acute (LBT Treatment) | 1 | 0 | 1 | 0 | 0 | 0 | 0 | 2 |
| 2B | Acute (LBT Treatment) | 1 | 0 | 1 | 0 | 0 | 0 | 0 | 2 |
| 2LL | Acute (LBT Treatment) | 2 | 0 | 1 | 1 | 1 | 0 | 1 | 6 |
| 3N | Acute (LBT Preventative) | 2 | 0 | 2 | 3 | 1 | 2 | 1 | 11 |
| 3L | Acute (LBT Preventative) | 3 | 0 | 2 | 4 | 2 | 3 | 2 | 16 |
| 3R | Acute (LBT Preventative) | 3 | 0 | 1 | 4 | 2 | 3 | 2 | 15 |
| 3B | Acute (LBT Preventative) | 3 | 0 | 4 | 4 | 2 | 3 | 2 | 18 |
| 3LL | Acute (LBT Preventative) | 3 | 1 | 2 | 3 | 2 | 2 | 2 | 15 |
